# Colour polymorphism and conspicuousness do not increase speciation rates in Lacertids

**DOI:** 10.1101/2023.02.15.528678

**Authors:** Thomas de Solan, Barry Sinervo, Philippe Geniez, Patrice David, Pierre-André Crochet

## Abstract

Conspicuous body colours and colour polymorphism have been hypothesized to increase rates of speciation. Conspicuous colours are evolutionary labile, and often involved in intraspecific sexual signalling and thus may provide a raw material from which reproductive isolation can easily evolve, while polymorphism could favour rapid evolution of new lineages through morphic speciation. Here, we investigated the influence of the presence/absence of conspicuous colourations, and of colour polymorphism on the speciation of Lacertids. We used several state-dependent diversification models, and showed that, regardless of the methods, conspicuous colourations and colour polymorphism were not related to species speciation. While the lack of correlation between conspicuous colourations and speciation rates is in line with most of the literature testing this hypothesis, the results for colour polymorphism contradict previous studies, and question the generality of the morphic speciation hypothesis.

## INTRODUCTION

Species diversification is not a constant process across time and taxonomic groups (Rabosky et al. 2007; Diaz et al. 2019). Within cephalopods for example, the Octopus order contains hundreds of species, while the much older order Nautilida only contains three species (Lindgren et al. 2004). Numerous studies have tried to explain these variations in diversification rates by investigating the links between ecology, life history traits or phenotype and extinction or speciation rates (Cardillo et al. 2003; Arbuckle and Speed 2015; Cooney and Thomas 2020). Among the traits investigated, those under sexual selection (sexual traits) have attracted a lot of attention because divergence in sexual traits can generate reproductive isolation between parapatric (Boul et al. 2007) or allopatric lineages (Price 1998; Panhuis et al. 2001) and strengthen reproductive barriers during secondary contact (Svedin et al. 2008), hence increasing speciation rates. However, sexual selection may also impose a burden on population viability and hence to increase extinction rates (Houle and Kondrashov 2002; Doherty et al. 2003; Kokko and Brooks 2003). The outcome of these opposing processes is difficult to predict and, until now, studies linking sexual selection and diversification rates have produced mixed results (Barraclough et al. 1995; Seehausen 2000; Kraaijeveld et al. 2011; Huang and Rabosky 2014; Cooney et al. 2017).

Another trait long suspected to increase the rates of speciation is polymorphism, here defined as the co-occurrence of several discrete, heritable morphs co-occurring within populations. Polymorphic species have been suggested to have higher speciation rates, based on two lines of reasoning resting mainly on indirect or theoretical arguments.

The first line of reasoning is that polymorphism may increase the rate of speciation if morphs are associated to different ecological niches (i.e. ecomorphs). The presence of alternative ecomorphs within the same population widens the ecological niche of the species. This may in turn facilitate range expansion through an increase of colonization abilities and the persistence of populations in variable environments (Forsman et al. 2008; Takahashi and Noriyuki 2019), two factors that are known to increase speciation rates (Kennedy et al. 2017). Specialized ecomorphs have been observed in several species, but it is not yet clear whether they occur in lacertids. (Kusche et al. 2015; Lattanzio and Miles 2016; Whitney et al. 2018, Scali et al. 2016).

The second line of reasoning rests on the “morphic speciation” scenario (West-Eberhard 1986). According to this scenario, a polymorphic population can lose a morph during the colonization of a new environment or area, resulting in a rapid phenotypic divergence of the remaining morph(s) and ultimately in reproductive isolation from the ancestral population (Corl et al. 2010*a*). The rapid phenotypic divergence after the loss of one morph may occur through (i) the cessation of gene flow from the missing morph, which used to continuously reintroduce maladapted alleles into the other morphs’ genetic backgrounds, preventing them from reaching their phenotypic optima (character release, West-Eberhard 1986); (ii) a change in the fitness landscape, as optimal trait combinations for one morph may change when another morph disappears from the population. For example, loss of a morph can drastically change the predation pressure acting on the remaining morphs (Bond 2007) if predation generates frequency dependent selection (Olendorf et al. 2006). Similarly, competition among morphs can generate frequency-dependent selection in polymorphic species (Sinervo and Lively 1996).

However, other lines of arguments suggest that polymorphism could reduce speciation rates and that the effect of polymorphism depend on mate choice within polymorphic populations. While assortative mating by morph could indeed enhance speciation and promote shifts from ancestral polymorphism to derived monomorphism through speciation (Jamie and Meier 2020), disassortative mating by morph is likely to slow speciation and retain polymorphism through the speciation process (Jamie and Meier 2020). As reported in *Heliconius numata* (Chouteau et al. 2017), disassortative mate preferences based on color morph has the potential to hamper ecological specialization by enhancing the homogenization of genomic backgrounds; ultimately preventing ecological speciation. Last, we argue that random mating relative to morph should promote less stringent mate choice in polymorphic species than in monomorphic species, as preferences span a larger spectrum of phenotypic space than in monomorphic species. In such cases (random or disassortative mating), the changes in colour traits should be less likely to result in reproductive isolation compared to non-polymorphic species. To sum up, the effect of colour polymorphism on speciation rate could depend on mate choice within population, which casts doubt on the generality of the morphic speciation hypothesis.

In spite of the theoretical interest generated by the “morphic speciation” hypothesis, this hypothesis has been examined empirically by only four studies until now. First, two studies investigated the effect of polymorphism on population divergence at small evolutionary scales (within a species complex in lizards, Corl et al. 2010*b*, 2012). They found that polymorphic populations were often ancestral, and that populations that lost a morph after colonizing a new environment showed reproductive incompatibilities with the ancestral population and rapid phenotypic changes. At macroevolutionary scale, two studies found that polymorphism increased speciation rates in birds and in lacertid lizards (Hugall and Stuart-Fox 2012; Brock et al. 2021). Taken together, these findings support the hypothesis of increased speciation rates in polymorphic species.

The lacertid lizards (family Lacertidae) is a group of Squamata containing more than 300 species distributed across Europe, Africa and Asia. This family is split in two main clades, the Gallotiinae, which contains only a few and often insular species, and the Lacertinae, which contains most of the lacertids species. The Lacertinae are divided between (i) the tribe Eremiadini, that lives mainly in xeric habitats in North African and the Middle East and often display dull colourations, and (ii) the Lacertini, that live mostly in temperate habitats in Europe and are more often colourful. Among those three clades, many species present conspicuous colourations. Bright colourations are found in both sexes on the throat and the belly, and are known to influence female mate choice and male-males contests (Abalos et al. 2016; Badiane et al. 2020). In some species, males also display blue or green ocelli on their flanks, or conspicuous outer ventral scales, that serve as indicators of the male quality during males intrasexual competition (Pérez i De Lanuza et al. 2014; Names et al. 2019). Furthermore, several lacertids present a polymorphism of colour. Such polymorphism can be found in both sexes when it occurs on ventral colouration, but is restricted to males when it occurs on the flank. In the genus *Podarcis*, the Common Wall Lizard *P. muralis* has become a model species for the study of colour polymorphism as several morphs differing in ventral colouration coexist in both sexes in most populations (Galeotti et al. 2013). These morphs may be related to different breeding strategies and life history traits (Calsbeek et al. 2010; Galeotti et al. 2013; but see Abalos et al. 2020) and they have a simple genetic determinism in *P. muralis* and in six other congeneric species, being controlled by two small regulatory genomic regions (SPR and BCO2, Andrade et al. 2019). Overall, the presence of specialized morphs and the variability of sexual colourations make the family Lacertidae an ideal model to investigate the impact of sexual selection on speciation rates and the hypothesis of morphic speciation.

Here, we use colouration data obtained from literature and photographs of live specimens for most species of the family Lacertidae (see “Data acquisition” for species excluded and source of information) to address two questions: (i) what are the evolutionary histories of conspicuous colourations and colour polymorphism in this family? (ii) Is there an effect of conspicuousness and colour polymorphism on speciation rate? We hypothesized that brightly coloured species undergo higher intensity of sexual selection than dull ones. However, because several previous studies failed to find a link between sexual selection proxies and speciation rates (Huang and Rabosky 2014; Cooney et al. 2017), we did not expect an effect of conspicuous colourations on speciation. For colour polymorphism, based on the morphic speciation hypothesis, we predicted that colour polymorphism increases speciation rates in lacertids. During the course of our study, we became aware of the publication of Brock et al. (2022) who had independently addressed the same question using the same group of species but using different data and methods (see discussion). Although our study started out as a completely independent work, unknowing that another group was addressing the same subject, it is important to understand why we came to opposite conclusions to Brock et al. (2022) and have devoted a section to address the reasons that might explain this discrepancy.

## MATERIAL AND METHODS

### Data acquisition

We follow the taxonomy and species list of lacertids from the Reptile Database (Uetz et al. 2020). We removed from this list several categories of species that are expected to have speciation modes differing from the rest of the species and driven by mechanisms presumably not be affected by the presence of polymorphism. We first removed the parthenogenetic species, as they arise by interspecific hybridization and result in unisexual clones providing “instant” reproductive isolation from their parent species (several species of the genus *Darevskia*, noted as “parthenogenetic” in Table S1). We also removed strictly insular species (species noted as “insular” in Table S1). Although morphic speciation might have helped the divergence of some insular species, we believe that geographic isolation remains the primary factors in of the speciation process. Supporting this idea, no case of in situ (intra-island) speciation is known in insular lacertids. We also removed four species for which we could not find any accurate information on colouration and five species with an uncertain taxonomic status (as judged by two of us, PAC and PG). Insularity was assessed using the distribution information available in the reptile-database website. After these steps, we retained 295 species for the speciation analysis. The final list of all species, including all species removed from the dataset, can be found in Supplementary Table S1.

We scored the presence of sexual colourations and polymorphism from multiple sources, such as scientific and naturalist papers and field guides, but also multiple photographs taken by the authors in the field (sources listed in Table S1). A species was considered as having conspicuous colourations if, at least in males, (1) the ventral side, or a part of its ventral side (throat, belly), was not white or grey, or (2) the flanks displayed several ocelli or spots that contrast with the rest of the dorsal/flank areas and are of a different colour, or (3) the species displayed a specific colour during the mating season. Furthermore, we considered a species as polymorphic if, within a population, individuals of a same sex and age-class exhibit several clearly distinct colours on the same body region. We did not score colouration as polymorphic if the morphs were not clearly distinguished, nor if the distinct morphs were associated with age or sex, or correspond to different parts of the annual cycle (green dorsal coloration in males of many species of Lacertinae disappear outside breeding season). Species where colouration varies geographically but not within populations were not treated as polymorphic, even though it was sometimes difficult to determine if some variation in colouration reported in the literature was geographically structured or not. Finally, a few species display polymorphism on dorsal pattern that is not linked to conspicuous coloration (different types of patterning of brown female dorsal coloration in *Podarcis* for example, see Ortega et al. 2014; Ortega et al. 2015). Such polymorphism is rare and always occurs in species which also show polymorphism for conspicuous ventral coloration; as a consequence, there is no species with polymorphism but not conspicuous coloration in our data. Data on sexual colourations and polymorphism are provided in Supplementary Table S1and some examples are provided in Figure S1.

We used the phylogeny provided by Garcia-Porta et al. (2019) as backbone phylogeny. We subsequently added the 85 non-insular and non-parthenogenetic species that were not sampled by Garcia-Porta. These species were added to the backbone tree using the function add.species.to.genus from the phytools package (Revell 2012). For 17 species (listed in Table S1) where the phylogenetic position was known from previous works, the species was added with its sister taxa at a random divergence date. If no phylogenetic information was available, the species was randomly located in its genus; this applied to 68 species in our phylogeny, of which only four were treated as polymorphic (see Table S1). We repeated this operation one hundred times in order to account for phylogenetic uncertainty generated by randomly added species in all analyses.

### Trait evolution and ancestral state reconstruction

We investigated the evolutionary inertia for conspicuous colourations and polymorphism using the δ values (Borges et al. 2019). This index is designed to measure phylogenetic signal in categorical traits, with high δ value indicating high phylogenetic signal (i.e. strong phylogenetic constraint to trait evolution). To test the significance of the observed δ, we compared with a Wilcoxon test the distribution of the 100 observed δ, measured on the 100 phylogenetic trees, against a null distribution obtained by measuring δ after randomization of the trait data among species (phylogenetic signal = zero). In addition, we reconstructed the ancestral state of the colouration with the make.simmap function from the phytools package (Revell 2012). This function simulates stochastic character histories using the state of the character on the tips and a continuous-time reversible Markov model. For this analysis, colouration was coded with three states: (1) no conspicuous colouration, (2) presence of conspicuous colourations without polymorphism, (3) presence of conspicuous colourations and polymorphism. Transition rates between states were allowed to differ and we did not exclude transitions from (1) to (3) although no such transitions were recovered by the model (see below). We repeated the ancestral state reconstruction (ASR) on the 100 phylogenetic trees, with 100 simulations for each analysis.

### Character associated diversification analysis

We used several methods to compare the rates of speciation between species with and without conspicuous colours, and between species with and without polymorphism. We excluded the species lacking conspicuous colourations of the polymorphism analysis because there are no direct evolutionary transitions from non-conspicuous to polymorphic, see results of the ASR). We however tested the influence of this exclusion by repeating the polymorphism analysis with all the species.

First, we used the speciation/extinction analysis of BAMM (Rabosky 2014) to detect shifts in speciation rates along the trees. This analysis allows the speciation rates to vary in time and among branches and does not consider character states. We then applied the STructured RAte Permutations on Phylogenies analysis (STRAPP, Rabosky and Huang 2016) to test if the speciation rates measured with BAMM were different between species with and without conspicuous colourations and colour polymorphism. STRAPP compares the Mann–Whitney U-test statistic measuring the relationship between the binary character state and speciation rates against a null distribution.

Secondly, we inferred the speciation rates with the Diversification Rates statistic (DR statistic, Jetz et al. 2012). Despite its name, the DR statistic provides a better estimate of the rate of speciation than of net diversification (Title and Rabosky 2019). For a given species, the DR statistic is computed as a weighted average of the inverse branch lengths connecting the focal species to the root of the phylogeny. We then used a non-parametric test, FiSSE (Fast, intuitive State-dependent Speciation Extinction analysis, Rabosky and Goldberg 2017), to investigate the effect of conspicuous colourations and polymorphism on speciation rates. For a given tree, this test first measures the difference of mean DR statistic between species with different character states, and then compares this difference to a null distribution. When testing for polymorphism effect, we first measured the DR statistic on the whole tree, then removed the species lacking conspicuous colourations and performed the test.

Finally, we fitted State Dependent Diversification models (SDD) to test the influence of conspicuous colourations and colour polymorphism on the diversification rates of Lacertid. For each trait, we implemented four SDD models: two models measuring character dependent diversification, plus the corresponding two null models estimating Character Independent Diversification (CID). The first SDD model we fitted was a Binary-State Speciation and Extinction model (BiSSE, Maddison et al. 2007). This model estimates one rate of speciation and extinction for each character state. The second SDD model was the Hidden State Speciation and Extinction model (HiSSE; Beaulieu and O’Meara 2016). This model includes hidden states, which allows the diversification rates to vary within each state (e.g. polymorphic species can have two different diversification rates). We also fitted two Character Independent Diversification models: CID-2 and CID-4, which were the null models corresponding respectively to BiSSE and Hisse: (Beaulieu and O’Meara 2016). They respectively have the same number of parameters than BiSSE and HiSSE models, but differ in having their diversification parameters independent of the observed character state (i.e. diversification parameter set equal for the observed state and different for the hidden states). It should be noted that the diversification rates estimated by SDD and CID models are not time dependent, and only vary depending on characters states (unlike speciation/extinction analysis of BAMM; Rabosky 2014). We selected the best-fit model among these four models based on the Akaike Information Criterion corrected for sample size (AICc, models were preferred when delta AICc > 2). All the models were made using the R package Hisse (Beaulieu 2017).

To account for phylogenetic uncertainty, we fitted the four SDD models, as well as STRAPPS and FiSSE analysis for each trait and phylogenetic tree. For SDD models, we assessed the best-fitting model each time.

### Comparison with Brock et al. (2021)

Although our study and the study of Brock et al. (2021) were made on the same taxonomic groups, our analysis reached an opposite conclusion to the study of Brock et al. (see below). We thus made additional analyses to understand the origin of these differences. In order to explain the differences of results between our study and Brock et al.’s (2021), we performed the four SDD models included in main analysis, HiSSE, BiSSE, CID2 and CID4, as well as the STRAPPS and FiSSE analyses, while changing the factors that changed between the two studies.

i. Our analysis included more species than in Brock et al.’s analysis (294 vs 262), and we excluded insular species, which was not done in Brock et al.’s analysis. We tested how this difference of species sampling affected our results by repeating our analysis using only the species included by Brock et al. (2021). For this analysis we used species coloration character states as coded in our database and the R script as in our main analysis.
ii. Our coding of character states differed from Brock et al. (2021)’s in some species. Among the species we had in common, 10 species were coded as polymorphic in our dataset while they were coded as monomorphic in Brock et al. dataset, and 7 species were coded as monomorphic in our dataset while they were coded as polymorphic in Brock et al. dataset (see Table S2 for coloration data of Brock et al., and the rational of our coding). We tested how these differences affected our results by repeating the analysis with both the species and the character states used by Brock et al. (2021). The R script used for this analysis was the same as for our main analysis.
iii. Although we used the same R package (HiSSE; Beaulieu 2017), the HiSSE package was recently updated, and our analysis was based a more recent version than Brock et al. (2021). As a result, there are several differences in the R function between Brock et al.’s analysis and our analysis. This results in differences of functions (the function specifically built for CID4 models was removed in the recent version), but also in differences in the number of transition rates between states in SDD models. We tested how differences in R script affected our results by re-running the original Brock et al.’s analysis using their script, species and character states (hence repeating their analysis entirely without changing anything).

All the analyses were made on the phylogenetic tree produced by Garcia-Porta et al. (2019), which was used in both our main analysis and Brock et al. (2021). Brock et al. (2021) built a second phylogeny on which they repeated all their analysis, without affecting their conclusion. We did not used this phylogeny in the comparison because it was built with only five genes, while Garcia-Porta et al.’ phylogeny was built with one dataset of 324 anonymous nuclear loci and one dataset of 6269 protein-coding nuclear loci. In addition, some clades identified in Brock et al.’ phylogeny were at odd with previous results: for example, the Gallotiinae are nested within Lacertinae in Brock et al (2021), while they are sister clades in Garcia-Porta et al. (2019), Arnold et al. (2007) and Kapli et al. (2011).

## RESULTS

Conspicuous colourations were frequent in lacertids, as they were present in 63% of the species analyzed (Figure 1). Colour polymorphism, on the other hand, was rarer and only concerns 13% of the species analysed; it concerned the ventral region (throat and/or belly) in all polymorphic species except for the four species of the genus *Tropidosaura* where we identified polymorphism in the bright coloration of the lower flanks (see Tables S1, S2). Polymorphism was unequally distributed in lacertids: the subfamily Lacertini includes more than half of the polymorphic species while it only contained 100 species out of the 295 species of Lacertidae and within the Lacertini, all mainland species of the genus *Podarcis* were polymorphic species (see Table S1). Both traits showed high phylogenetic conservatism (median δ across phylogenies = 6.39, W=10e5, p<0.001 and median δ=75.7, W=10e5, p<0.001 respectively). The ancestral state reconstruction suggested more acquisitions than losses in the evolution of both conspicuous colourations and colour polymorphism (Figure S2, 43 acquisitions vs 23 losses for colourations and 15 acquisitions vs 3 losses for polymorphism). Transitions from dull-coloured to polymorphism, and the reverse, were very rare (0.04 and 1.1 on average across all trees respectively), indicating that, the polymorphic state evolves from a monomorphic conspicuous-colour state in lacertids. Finally, the ancestor of lacertids probably displayed conspicuous colourations without being polymorphic (Figure S2).

**Figure 1.**
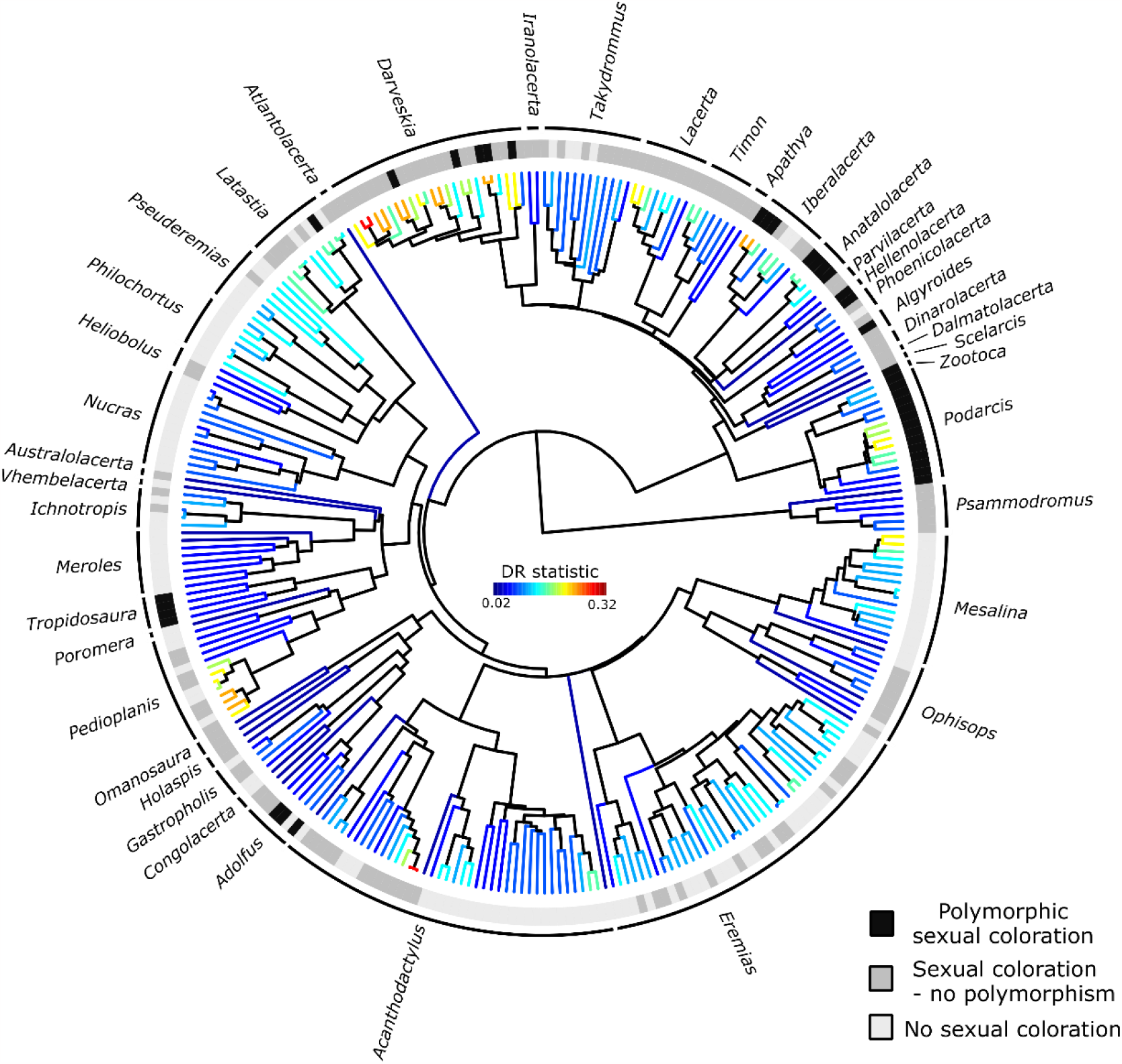
Phylogenetic relationships of the 295 Lacertids species included in the analysis. This tree was randomly chosen among the hundred trees produced for the analysis to account for phylogenetic uncertainty (see methods). States of sexual colouration and polymorphism characters are indicated at the tips. Tip branches are coloured according to the mean DR statistic measured across all trees (an estimate of the branch-specific speciation rate).

### Character associated diversification analysis

We wanted to test whether conspicuous colouration on the one hand, and polymorphism on the other hand, influence speciation rates. As explained above, to test for the effect of polymorphism, we excluded inconspicuous species, as the ancestral state of polymorphic lineages is always conspicuously coloured. Including inconspicuous species would confound the effects of polymorphism per se and conspicuousness. When included, however, species lacking conspicuous colourations had a minor impact on the results, and did not change the overall conclusion (see Supplementary Methods).

The BAMM analysis detected a shift of speciation rates only in 21 trees out of the hundred trees. In addition, the STRAPP tests indicated that neither the presence of conspicuous colourations nor the presence of colour polymorphism affected speciation rates (p>0.5 for all tests across the 100 trees).

The DR statistic showed that inconspicuously and conspicuously coloured species had similar speciation rates (average λ_0_ across trees = 0.11±0.04 and average λ_1_ across trees = 0.13±0.07 respectively, non-significant: p>0.5 for all FiSSE tests, Figure 1 and 2). Similarly, there was no difference in DR statistic between species conspicuously coloured monomorphic and polymorphic species (average λ_0_ across trees = 0.12±0.07 and average λ_1_ across trees = 0.14±0.07 respectively, non-significant: p>0.5 for all FiSSE tests, Figure 1 and 2).

**Figure 2.**
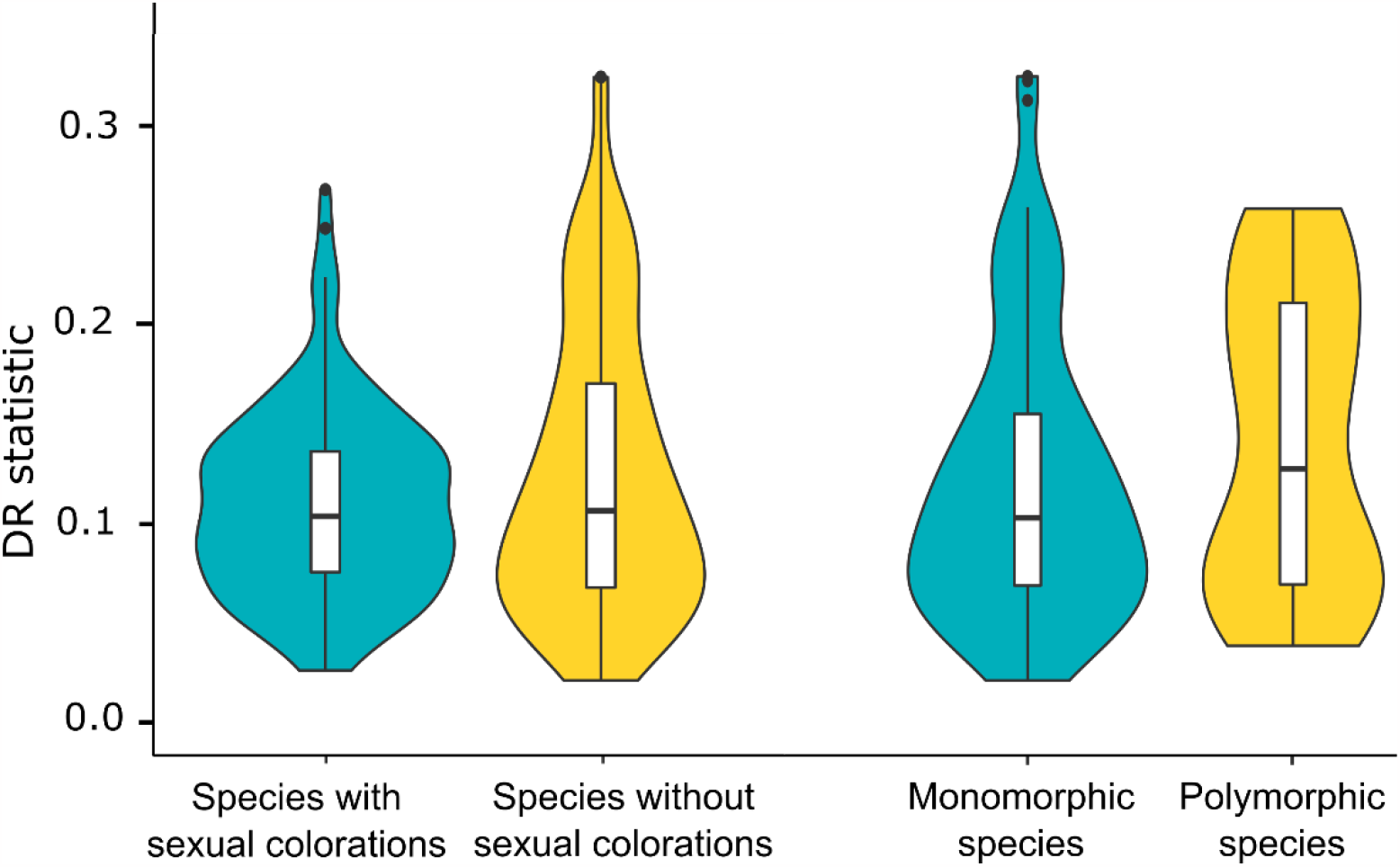
Violin plots of the DR statistic values for the species with and without sexual colouration, and with and without colour polymorphism.

Last, the best fitting State Dependent Diversification (SDD) models for the evolution of conspicuous colouration were models where diversification parameters are independent of the observed character state (CID 2 or CID 4) for all the trees (Figure 3). Similarly, we found a low support for an influence of colour polymorphism on the diversification of species with conspicuous colourations. CID 2 and CID 4 were the best fitting models for 75 and 22 trees respectively, while HiSSE was the best fitting model for 3 trees only (Figure 3).

**Figure 3.**
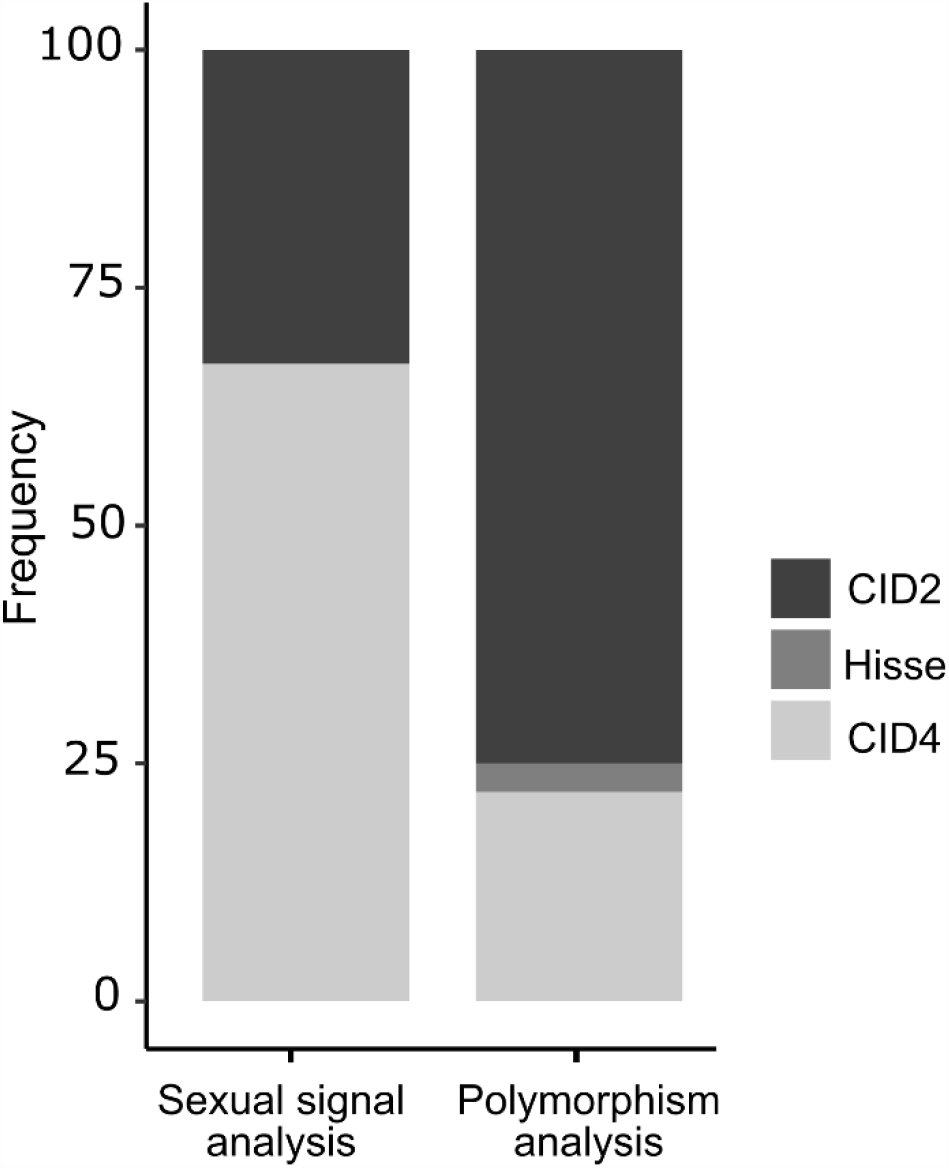
Frequency of the best fitting diversification models for the sexual colouration and colour polymorphism analysis repeated across one hundred trees. Models with character dependent diversification (HISSE) were considered as best fitting when delta AICc > 2. BiSSE models were included in both analyses, but never had the lowest AICc.

### Polymorphism analysis applied to all species, including those without sexual colorations

The results of STRAPP and FiSSE analysis were not affected by the inclusion of species without sexual colorations. Both analyses indicated that speciation rates were independent of the presence of color polymorphism (p>0.5 for all tests across the 100 trees).

When species without sexual colorations were included, SDD models indicated that color polymorphism was probably independent of the rates of diversification. The HiSSE model was the best-fitting model for only 2 trees, while CID2 and CID4 were the best fitting models for 44 and 54 trees respectively.

### Comparison with Brock et al. (2021)

The comparison with Brock et al.’s (2021) analysis indicated that the differences of results between our SDD models and theirs stem from differences in character states, i.e. in the species considered to be polymorphic *versus* not (for more details see Table S2), as well as from the use of an old version of the package encoding SDD models in Brock et al (2021). However, this difference of results only applied to SDD models. Unlike SDD models, the non-parametric FiSSE and STAPPS analysis never detected an effect of polymorphism on speciation in the set of species studied by Brock et al., whether we used our coding of character states (STRAPPS test: p=0.98; FiSSE test: λ_0_ = 0.10, λ_1_=0.13, p=0.88), or theirs (STRAPPS test: p=0.98; FiSSE test: λ_0_ = 0.10, λ_1_ = 0.14, p=0.98).

When we repeated our analysis with the same species as in Brock et al. (2021), the model with the lowest AIC was still CID4, like in our main analysis (AIC and parameters values for all SDD models in Table S3). This indicates that the inclusion of species not included in Brock et al. (2021) and the exclusion of insular species in our analysis were not at the origin of the difference of results.

When we built the SDD models with the species and character states of Brock et al. (2021), the results were opposite to what we previously found: the HiSSE model scored the lowest AIC, just before CID4 model (AIC_HiSSE_ = 1838, AIC_CID4_ =1841; see Table S3 for the score of the other models). The difference of SDD model’s results between our study and Brock et al. study thus lies in in the way of coding polymorphism.

When we used the R script, the species and the character states of Brock et al. (2021), we also found that the HiSSE model scored the lowest AIC (AIC_HiSSE_ = 1835), while the other SDD models all had much higher AIC values (AIC_BiSSE_ = 1862, AIC_CID4_ =1901, AIC_CID4_ =1913; parameters values in Table S3). Note that, compared to the recent version of the R package on the same data, the AICs obtained with the older version were lower for the state-dependent models (HISSE and BISSE) and much higher for non-state dependent models (CID2 and CID4). Note also that the SDD models did not perform properly in any analysis using Brock et al.’s data and character states, as 6 of the 8 estimated parameters were at the boundary of the allowed interval in our HISSE model (all four extinction rates and two transition rates are zero, see Table S3).

## DISCUSSION

Our analyses suggested that neither the presence of conspicuous colourations nor colour polymorphism increases the rates of speciation in Lacertids. We also showed that conspicuous colourations and colour polymorphism are labile across taxonomy, and were gained and lost several times during the diversification of Lacertids. The lack of effect of conspicuous colourations on speciation is in agreement with several previous studies (Huang and Rabosky 2014; Gomes et al. 2016; Cooney et al. 2017). However, our results for colour polymorphism is in contradiction with the two studies that tested the morphic speciation hypothesis at multispecies level (Hugall and Stuart-Fox 2012; Brock et al. 2021), including one that used the same family as model (Brock et al. 2021).

### Inconsistent results on Lacertids

The lack of effect of colour polymorphism on speciation rates contradicts the conclusions of Brock et al. (2021) obtained on the same family Lacertidae. Although Brock et al. (2021) used a different phylogeny, they also used the same phylogeny as us and this did not affect their conclusion.

We also excluded insular species and parthenogenetic species, because speciation mechanisms in these species are unlikely to depend on polymorphism, unlike Brock et al. (2021). In addition, their phylogeny only included species that were represented in the molecular dataset. While this avoided placing species randomly within their genus, as we did (we checked that our results were robust to the random position of species with no genetic data within their genus by repeating all analyses on 100 trees, see methods), this also resulted in removing a considerable fraction of the Lacertidae species from their data. Worryingly, this fraction of species excluded from their analysis is biased between the two main clades of Lacertidae (Table S1): their dataset lacks 17 species of Lacertini out of 143 but 66 species of Eremiadini out of 203. Since polymorphic species are mostly Lacertini, it seems dangerous to underrepresent the mostly non-polymorphic Eremiadini when comparing the diversification rate of polymorphic and non-polymorphic species. If we wanted to maintain our choice to include as many species as possible in the dataset, switching from Garcia-Porta et al’s tree to Brock et al’s tree would only decrease the number of randomly added species from 68 to 57. Last, Garcia-Porta et al’s backbone is based on genomic data while Brock et al’s tree, based on a small amount of sequence data, has obtained some spurious relationships. Finally, when we tested how this difference of species sampling affected our results by repeating our analysis using only the species included by Brock et al. (2021) and using species coloration character states as coded in our database, we found the same conclusion of no effect of polymorphism on diversification. We are thus confident that our choice of using Garcia-Porta et al.’s (2019) phylogeny and including species that have no genetic data available was the best methodological choice, but the use of different phylogenies and different species did not cause the differences in the two studies.

Another possible reason is that our attributions of character states (polymorphic versus nonpolymorphic) differed in some cases from Brock et al. (17 species highlighted in yellow in Table S2), and indeed repeating our analysis with the species and character coding of Brock et al. (but using our R script and phylogeny) suggested a possible influence of polymorphism on diversification, as the best model this time was the HiSSE model. We therefore re-checked our attributions in case of mismatches. Ten species considered monomorphic by Brock et al were considered polymorphic by us. Some of these reflect the fact that we coded polymorphism as the presence of discrete states on any conspicuous colouration, while Brock et al. (2021) considered only throat colouration -thus excluding four *Tropidosaura* species. For six other species we re-checked bibliographic references (see table S2) and confirmed mentions of polymorphism that had been apparently overlooked by Brock et al. 2021. Seven species were considered monomorphic by us and polymorphic by Brock et al. 2021. In four of these, colour morphs do not co-occur within a population, but occur in allopatric populations or subspecies – which does not qualify as polymorphism *sensu stricto*. For the last three, we did not find any mention of polymorphism in the literature (and Brock et al. 2021 did not provide a reference).

One of the reasons for the different conclusions between our study and Brock et al. is thus differences in species trait coding (see Table S2), partly resulting from differences in interpreting available evidence, partly resulting from a restriction of polymorphism to the throat coloration in Brock et al. All in all, we believe that, although the status of each particular species can always been updated and discussed, our character states are more accurate and closer to the original definition of a polymorphic species and to the concept of morphic speciation (i.e. not focused only on throat colouration, West-Eberhard 1986, and excluding geographical races).

In addition, methodological issues can contribute to discrepancies in the results. The SDD methods have been criticized on the basis of high type-1 errors, i.e. they may detect differences in diversification rates even for neutral traits and are probably overparameterized as speciation and extinction can hardly be estimated independently (Rabosky and Goldberg 2015, 2017; Louca and Pennell 2020). This is illustrated by the fact that 6 of the 8 estimated parameters are at the boundary of the allowed interval in the HISSE model (all four extinction rates and two transition rates are zero) when we used Brock et al.’s data and character states with the new version of HiSSE. The same problem was already apparent in Brock et al.’s original results (their Table 2, repeated here in Table S3). Especially worrying is the fact that the net turnover parameters are not estimated (parameters at boundary) for the two state classes (polymorphic and non-polymorphic) of the A class of the hidden state. The higher diversification of polymorphic species is thus only supported in one of the two classes of the hidden state using the new version of the R package HiSSE, which does not allow to conclude on the impact of polymorphism on diversification rates according to Beaulieu and O’Meara (2017). FiSSE and STRAPP have been developed to avoid this kind of drawbacks and provide a more robust, and nonparametric, way of testing character-dependent speciation rates (Rabosky and Huang 2016; Rabosky and Goldberg 2017). The fact that these methods do not recover an effect of polymorphism, whatever the data and character states used (ours or Brock et al’s), and that Brock et al. results are less robust when using their species and trait coding with the new version of HiSSE suggests that the result of Brock et al (2021) might have been partly (mostly?) driven by a type-1 error.

In conclusion, the differences between the results of Brock et al. (2021) and our results concerning the impact of colour polymorphism on diversification in Lacertidae originate partly from Brock et al.’s choice to restrain colour polymorphism to throat colour polymorphism only and partly from methodological issues associated with the R package HiSSE.

### Conspicuous colorations, sexual selection and speciation in lizards

The lack of effect of conspicuous colourations on speciation rates at the scale of a family contrasts with the accumulation of evidence showing an influence of sexual selection on pre-zygotic (Boughman 2001; Masta and Maddison 2002; Boul et al. 2007) and post-zygotic isolation (Vamosi and Schluter 1999; Naisbit et al. 2001) at short time scales. Yet, there is not necessarily a link between processes acting at short time scales and diversification rates observed at large phylogenetic scales. This decoupling could arise because different processes influence the creation of new species and their persistence over time. For instance, some authors found that sexual selection increases the risk of extinction in birds (McLain et al. 1995; Doherty et al. 2003; but see Cooney et al. 2017 for opposite results). However, SDD models were found to be unable to accurately measure both speciation and extinction rates (Beaulieu and O’Meara 2016; Louca and Pennell 2020), and this hypothesis is thus difficult to test using current comparative methods. The lack of correlation between speciation rates and the presence of sexual colourations could also come from the fact that speciation does not increase with the intensity of sexual selection but rather with the evolutionary rates of sexual traits (Cardoso and Mota 2008). These two features are not necessarily correlated, as a trait may be at the same time very conserved across species and under strong sexual selection (Song and Bucheli 2010; Mejías et al. 2020). The few studies investigating this question found support for a positive correlation between speciation rates and evolutionary rates of sexual traits (Cardoso and Mota 2008; Gomes et al. 2016) but they were made on birds, and more research on other taxonomic groups is needed. Finally, the occurrence of conspicuous colouration might be a poor proxy of the intensity of sexual selection. In lacertids, mating choice and interspecific recognition are known to rely on pheromones as much as on colouration (Cooper and Pèrez-Mellado 2002; Khannoon et al 2011; Gabirot et al. 2013). It is thus possible that considering the presence of conspicuous colourations, even when they are sexually dimorphic, is not sufficient to detect an effect of sexual selection on speciation.

An additional explanation for the lack of correlation between conspicuous colourations / polymorphism and speciation rates could be the limited role that pre-zygotic isolation has in lacertids diversification. The influence of sexual selection and polymorphism on species speciation occurs mainly at the pre-zygotic stage (West-Eberhard 1986; Coyne and Orr 1989; Gray and Cade 2000; Ritchie 2007). However, the influence of this stage in lacertids speciation is unclear: some species are partially able to recognize each other using pheromones (Barbosa et al. 2006; Gabirot et al. 2012), but interspecific courting seems to be frequent (Martín and López 2006; Galoyan et al. 2019), suggesting that post-zygotic isolation also play a strong role in the speciation process (Carretero 2008; Pinho et al. 2009). On the other hand, the groups in which an effect of sexual selection and polymorphism was reported are taxa where the pre-zygotic isolation is determinant for speciation (e.g birds, Barraclough et al. 1995; Brambilla et al. 2008). This may explain why, depending on the groups, sexual trait and polymorphism sometimes correlate with speciation rates and sometimes do not. We also lack data on the effect of morphs on mate choice in Lacertidae, as we only found some information on mate choice relative to morphs in *Podarcis muralis* (Pérez i de Lanuza et al. 2013; Sacchi et al. 2015, 2018, who found mixed support for colour assortative mating) while indirect genetic results also suggest some possible assortative mating in *Podarcis melisellensis* (Huyghe et al. 2010).

One main limitation of our work lies in the data we used to score the colouration. Neither photographs nor literature sources account for variation of ultraviolet (UV) colouration. Lacertids frequently use UV colouration for mate choice or to signal their fighting capacity (Olsson et al. 2011; Pérez i de Lanuza et al. 2014). Thus, in principle, we could have underestimated the frequency of conspicuous colouration and polymorphism in our dataset. However, known UV signals in lizards are always displayed on a patch with a visibly different colour than the rest of the body (blue throat or green ocelli for instance), so it is unlikely that this issue affects our detection of conspicuousness, although it may have done so for polymorphism. As far as we know, there is currently no example of polymorphism limited only to UV colouration in animals. A second methodological limitation may stem from the strong geographical bias prevailing in the taxonomic information. In the last twenty years, an intensive work unraveled the phylogeny of the lacertids of the northern hemisphere, allowing the description of numerous species. On the other hand, almost no taxonomic studies based on genetic data have been made on lacertids of the equatorial region. The diversity of species living in this region is thus probably strongly underestimated. It is difficult to predict in which ways this issue may influence our results, because species with and without conspicuous colourations and colour polymorphism are present in the equatorial region. However, we can predict that undescribed species are more abundant in the Eremiadini (where colour polymorphism is rare) than Lacertini (where colour polymorphism is widespread, see Fig. 1 and Table S1), because taxonomic revisions for many Lacertini have already been published and undescribed cryptic diversity remains higher now in Eremiadini (pers. obs.). This should result in higher diversity and hence higher diversification rates for Eremiadini than currently estimated, and hence further reinforce the lack of difference between polymorphic and non-polymorphic species in terms of diversification rate.

A large variety of factors, besides those examined in this study, may explain the variations of speciation rates observed within groups (for instance diet, Tran 2016; habitat specialization Liedtke et al. 2016 or reproduction mode, Lynch 2009). In lacertids, the two most diverse genera (*Acanthodactylus* and *Eremias*, representing 22% of all the Lacertid species) live in arid areas of the Middle East or North Africa. Similarly, an increase of lizard diversity in arid areas have been reported in biogeographic studies (Powney et al. 2010; Lewin et al. 2016). As such, this pattern suggests that hot and arid conditions play a strong role in the radiation of lacertids. Yet, the factors at the origin of these radiations are still unclear. Adaptive radiation seems unlikely, as most lacertids species living in arid areas seem to show similar ecological niches as inferred from distribution patterns: most arid regions of the Middle East or North Africa are inhabited by a maximum of 3-4 species of *Acanthodactylus* (large bodied) and 3-4 species of *Mesalina* (small bodied) segregated by habitat preferences within genus (e.g. Haas 1951; Blanc 1980; Werner 1982; Schleich et al. 1996; Nouira and Blanc 2003; Rifai et al. 2003; Baha El Din 2007), and most species diversity in the genera correspond with changes in these species between regions. However, such pattern could be the result of the strong relation between the environment and ecological and life history traits of lizards (e.g. increase in activity time and fecundity with temperature; Adolph and Porter 1993) that may have facilitated speciation in arid areas (discussed in Powney et al. 2010). Nevertheless, currently no studies properly tested this possible link, and this hypothesis is yet to be confirmed.

## Conclusion

Our analyses suggest that the speciation of lacertids was not influenced by the presence of conspicuous sexual colourations nor by colour polymorphism. These results support the idea of a decoupling of the effect of sexual selection on species diversification between short time scale (i.e. effects on pre- and post-zygotic isolation) and large time scale (i.e. effects on rates of extinction/speciation). They also call into question the generality of the morphic speciation hypothesis, which is currently supported by only two large scale study (Hugall and Stuart-Fox 2012; Brock et al. 2021). The fact that prezygotic isolation does not appear to be determinant for the speciation in lacertids may explain this lack of effect of polymorphism and conspicuous colouration that we found. In the future, it would be interesting to test this hypothesis by investigating other taxa in which speciation is mainly driven by post-zygotic isolation.

We also wish to stress a previously neglected inherent difficulty with the detection of the morphic speciation model through comparative methods in empirical datasets. The diversification analyses used by us or by Brock et al. (2021) test for differences in diversification rates associated with the presence of polymorphism. Yet, the morphic species model predict that “polymorphic lineages should be ancestral and monomorphic lineages should be derived” (Corl et al. 2010b), as morphic speciation generates monomorphic species from polymorphic species. The signature of the morphic speciation model in phylogenies should thus be an excess of monomorphic species sister to a polymorphic species, not an increased diversification rate in clades where all species are polymorphic. This creates an overlooked paradox: detecting an increased net diversification rate in clades where most species are polymorphic would actually run against the predictions of the morphic species model, unless one assumes that some lineages have an inherent tendency to regain the polymorphic state quickly after it is lost – an assumption without clear support in the data.

## Acknowledgements

We thank all photographers who contributed to the amphibians and reptiles photo collection managed by PG.

## Data, scripts, code, and supplementary information availability

Data and scripts are available online: DOI: 10.5281/zenodo.7619485 (https://zenodo.org/record/7619485)

