## Supplementary Figures for "Colour polymorphism and conspicuousness do not increase speciation rates in Lacertids"

**A**

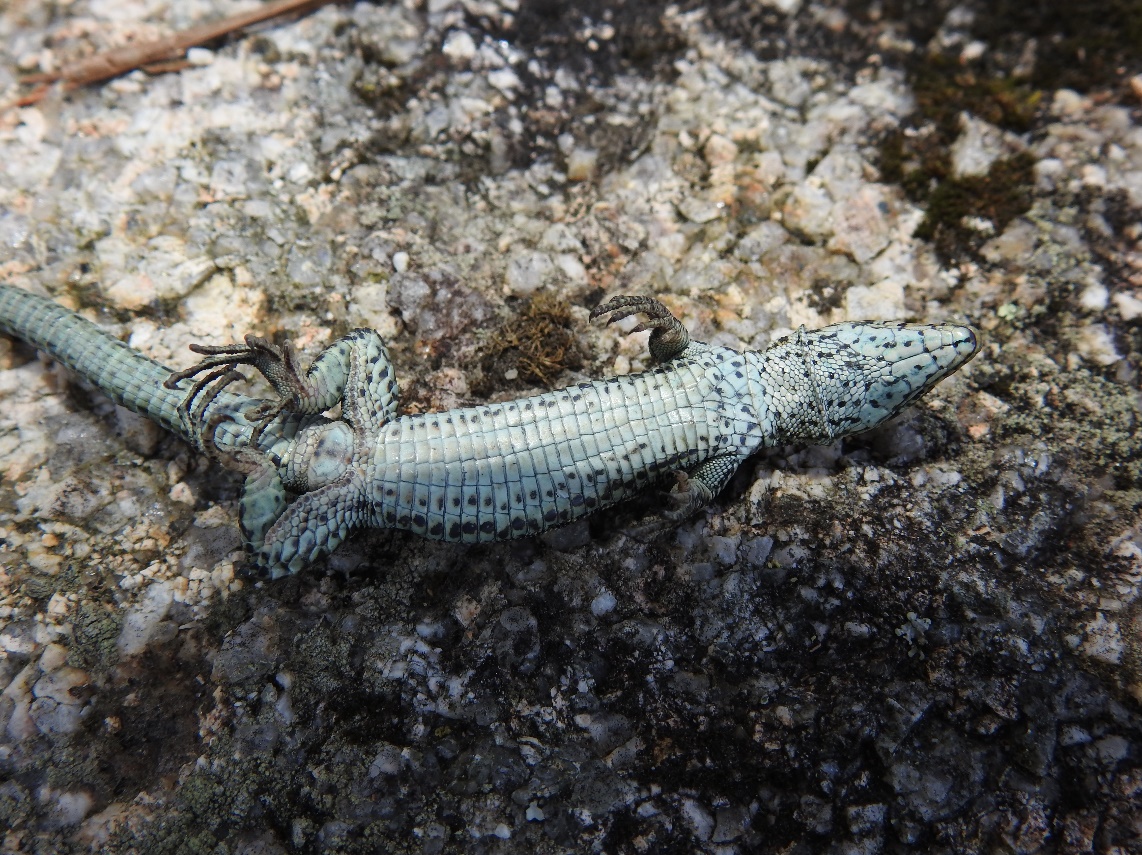

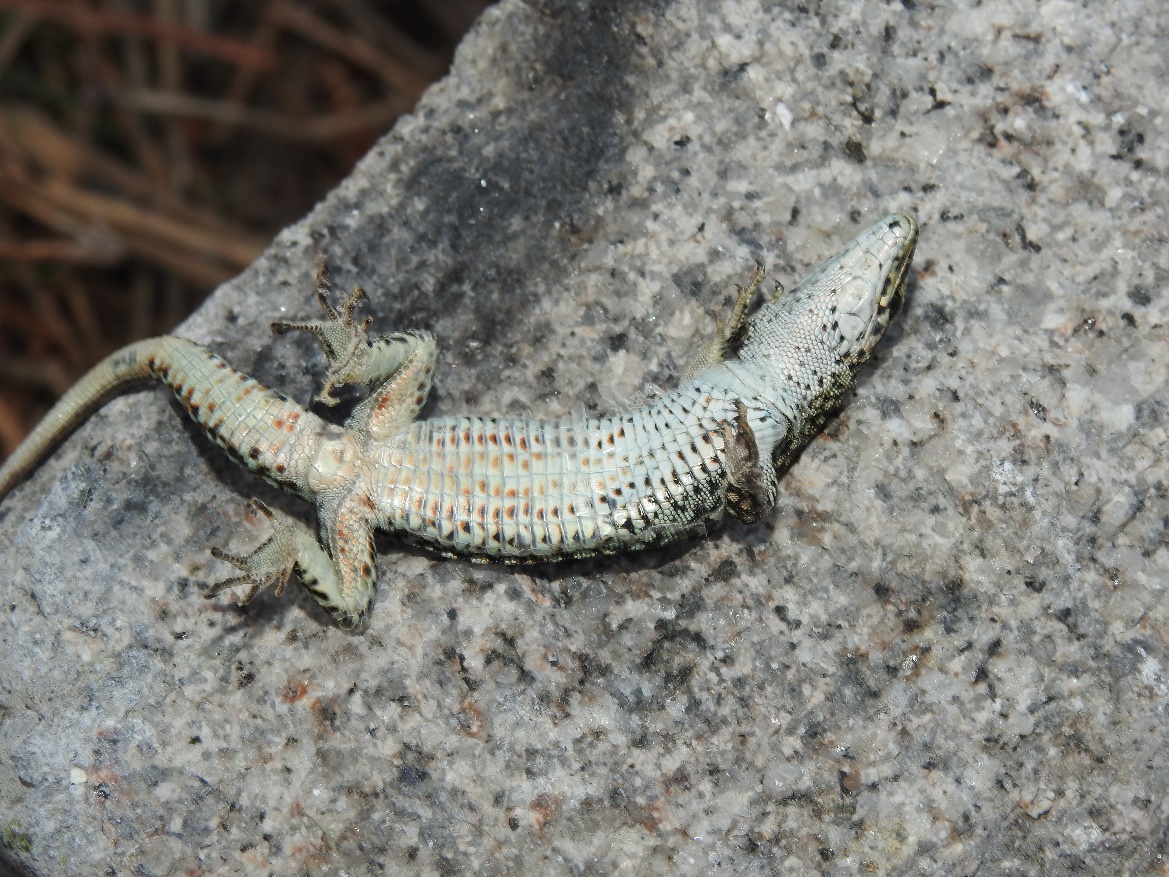

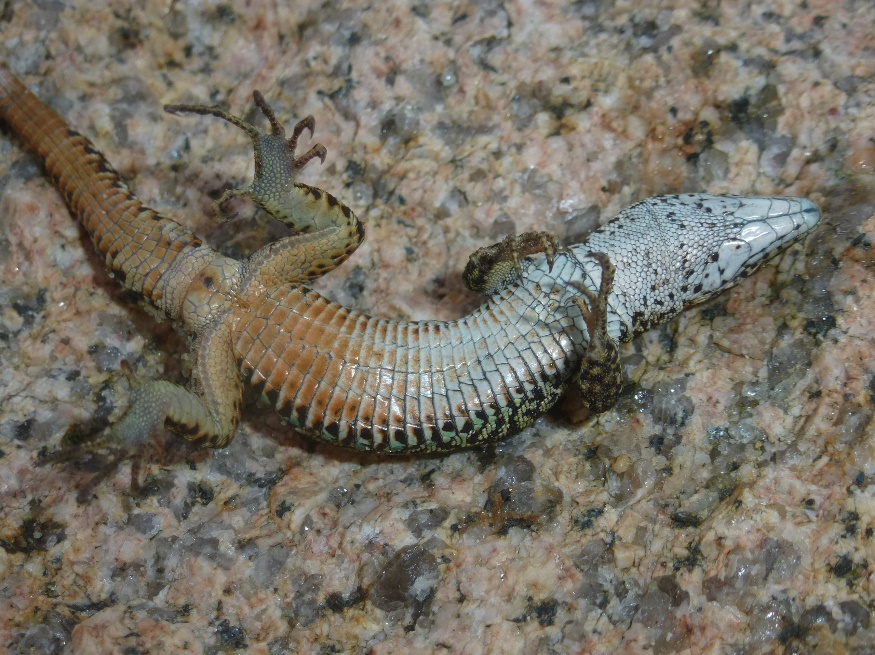

**B**

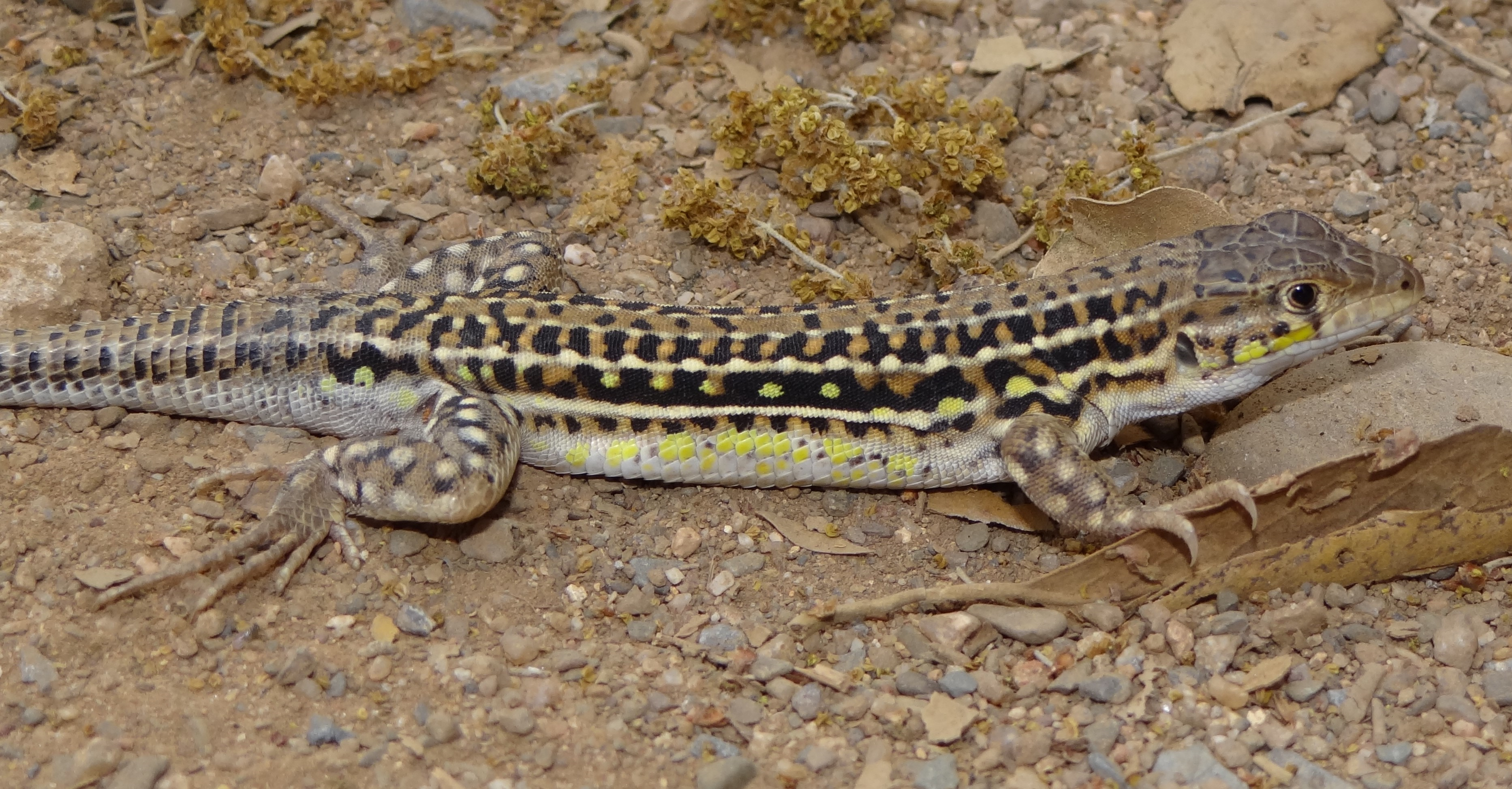

**Figure S1.** **A**: Polymorphism in conspicuous ventral coloration not associated with throat polymorphism in *Podarcis lusitanicus* (included as a subspecies within *P. guadarramae* in Brock et al. and in our study as the species was not recognised at the time). Three adult males from the same population (around O Pindo, Galicia, Spain) are depicted. **B**: Conspicuous coloration in an adult male *Acanthodactylus erythrurus* from near Bab Taza, Rif Mountains, Morocco.

All photos from PAC.

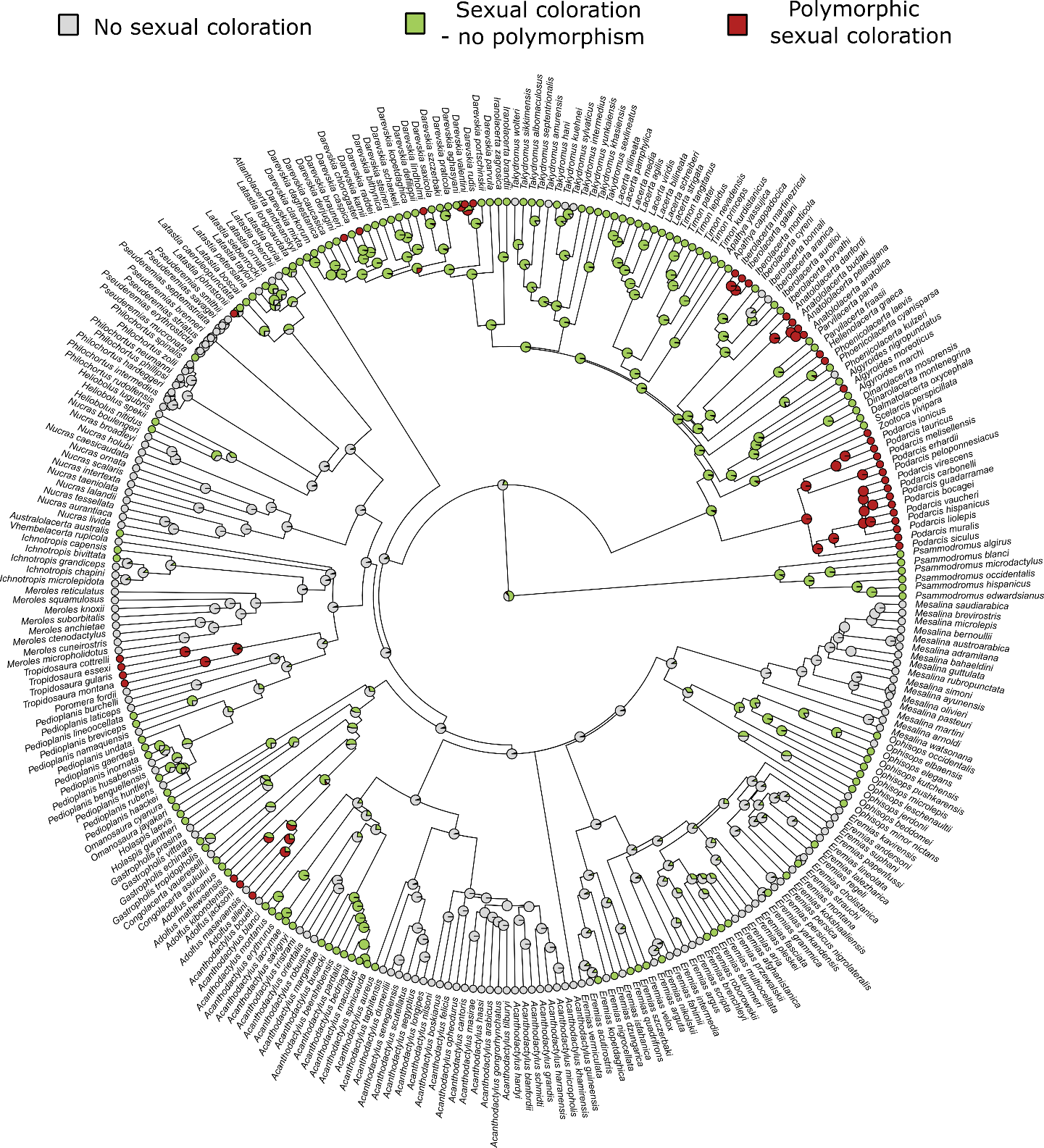
 **Figure S2.** Ancestral state reconstruction (ASR) of Lacertids coloration estimated for one of the 100 phylogenic trees. The ASR was made with the make.simmap function of the phytools package, using a continuous-time reversible Markov model. Transition rates were allowed to differ between states.
